# Neuronal subtypes and connectivity of the adult mouse paralaminar amygdala

**DOI:** 10.1101/2024.01.11.575250

**Authors:** David Saxon, Pia J Alderman, Shawn F Sorrells, Stefano Vicini, Joshua G Corbin

## Abstract

The paralaminar nucleus of the amygdala (PL) is comprised of neurons which exhibit delayed maturation. PL neurons are born during gestation but mature during adolescent ages, differentiating into excitatory neurons. The PL is prominent in the adult amygdala, contributing to its increased neuron number and relative size compared to childhood. However, the function of the PL is unknown, as the region has only recently begun to be characterized in detail. In this study, we investigated key defining features of the adult PL; the intrinsic morpho-electric properties of its neurons, and its input and output connectivity. We identify two subtypes of excitatory neurons in the PL based on unsupervised clustering of electrophysiological properties. These subtypes are defined by differential action potential firing properties and dendritic architecture, suggesting divergent functional roles. We further uncover major axonal inputs to the adult PL from the main olfactory network and basolateral amygdala. We also find that axonal outputs from the PL project reciprocally to major inputs, and to diverse targets including the amygdala, frontal cortex, hippocampus, hypothalamus, and brainstem. Thus, the adult PL is centrally placed to play a major role in the integration of olfactory sensory information, likely coordinating affective and autonomic behavioral responses to salient odor stimuli.

**Significance Statement:** Mammalian amygdala development includes a growth period from childhood to adulthood, believed to support emotional and social learning. This amygdala growth is partly due to the maturation of neurons during adolescence in the paralaminar amygdala. However, the functional properties of these neurons are unknown. In our recent studies, we characterized the paralaminar amygdala in the mouse. Here, we investigate the properties of the adult PL in the mouse, revealing the existence of two neuronal subtypes that may play distinct functional roles in the adult brain. We further reveal the brain-wide input and output connectivity of the PL, indicating that the PL combines olfactory cues for emotional processing and delivers information to regions associated with reward and autonomic states.

## Introduction

The paralaminar nucleus (PL) is a subregion of the mammalian amygdala containing neurons that are born prenatally but delay their maturation until adolescence (deCampo & Fudge, 2012; Sorrells et al., 2019). The late development of PL neurons contributes to a hallmark increase in human amygdala size and neuron number between childhood and adulthood (Avino et al., 2018). The PL was originally identified in the human (Crosby & Humphrey, 1941), and later in the monkey, sheep, tree shrew, and rat (Ai et al., 2021; Chareyron et al., 2012; Martí-Mengual et al., 2013; Nacher et al., 2002; Piumatti et al., 2018). We recently identified its homolog in the mouse, revealing similarities in its developmental and molecular features across the mouse and human (Alderman et al., 2023).

The mouse PL is located adjacent to the basolateral amygdala and ventral to the striatum. It is distinguished from nearby regions by its dense arrangement of small neurons that express markers of immaturity (Dcx, Psa-Ncam) and glutamatergic identity (VGlut1, VGlut2, Tbr1) during juvenile ages (P7-21). By early adulthood (P60), mouse PL neurons have matured (Dcx-, NeuN+), and most (∼84%) express the excitatory marker Tbr1. Thus, the PL is conserved across mammalian species as a nucleus of late-maturing excitatory neurons, and upon maturation it is likely integral to adult amygdala function. However, beyond histological analyses, the adult PL has not been well studied.

While the circuital and functional fate(s) of late-maturing neurons in the PL are unknown, late-maturing neurons have been studied in other brain regions, highlighting their significance to the adult brain (Benedetti et al., 2023). These regions include the rodent dentate gyrus and olfactory bulb, where the generation of newborn neurons occurs postnatally (Denoth-Lippuner & Jessberger, 2021; Sakamoto et al., 2014), as well as in the non-neurogenic piriform cortex, where prenatally-born neurons undergo gradual maturation similar to the PL (Rotheneichner et al., 2018). Across these regions, immature neurons complete their development into mature neurons with functional synaptic input (Benedetti et al., 2019; Dieni et al., 2019; Li et al., 2018). This integration is widely believed to contribute to activity-dependent region-specific learning (Benedetti & Couillard-Despres, 2022; Borzello et al., 2023; Chaker et al., 2023). Our previous studies have shown that the mouse PL develops a responsiveness to olfactory stimulation between juvenile and adult ages (Alderman et al., 2023). This suggests that like other late-maturing populations, PL neurons become functionally integrated upon maturation, and may be critical for adult behavior.

The identity of a neuron within a circuit is defined by both intrinsic and extrinsic characteristics. In addition to gene expression profiles, intrinsic characteristics include electrophysiological and morphological properties, which in many brain regions are tied to divergent roles (Ascoli et al., 2008; S. Chen et al., 2022; Gouwens et al., 2018; Hattox & Nelson, 2007; Sun et al., 2013). Extrinsic characteristics, defined by the regions and patterns of input and output connectivity, determine the functional domain of a brain region (Zeng, 2018), and are of particular interest in dissecting late-maturing populations (Benedetti & Couillard-Despres, 2022; Borzello et al., 2023). While our past studies have investigated the developing intrinsic properties in the PL at juvenile ages (Alderman et al., 2023), the electrophysiological profiles, morphologies, and connections that define the mature PL in adulthood are unknown.

Using a combination of electrophysiological, morphological, and anatomical tracing approaches, in this study we define the intrinsic neuronal properties of the adult PL as well as its input and output projections. We reveal that mature PL neurons are not uniform, but instead comprise two subtypes defined by different action potential firing patterns and dendritic branching properties. We further uncover bidirectional connections between the PL and the olfactory network and basolateral amygdala, with additional outputs to regions involved in memory, reward, visceral and autonomic states. Thus, the adult PL likely plays a central role in the integration of sensory information for appropriate behavioral processing.

## Methods

### Patch-Clamp Electrophysiology

Adult (P59-P78) C57/Bl6 mice (Jax #000664) (n = 12 males, 12 females) were anesthetized with isofluorane and decapitated. Brains were dissected and placed into ice-cold cutting ACSF, containing (in mM): Sucrose 234, D-Glucose 11, KCl 2.5, NaHCO3 26, NaH2PO4 1.25, MgSO4 7, CaCl2 0.5. Sagittal sections (275 µm thick) were sliced using a Leica V1200 vibratome. Sections containing the PL were transferred to recording ACSF, containing (in mM): NaCl 125, KCl 3.5, CaCl2 2, MgCl2 1, NaH2PO4 1.25, NaHCO3 26, D-Glucose 10. This solution was initially held at 34°C for 30 minutes to facilitate slice recovery, before cooling to room temperature for the duration of the experiment. All solutions were adjusted to 7.4 pH and 300 mOsm and were bubbled with carbogen (95% CO2/5% O2) during all steps. All chemicals were supplied by Sigma (St Louis MO)

Patch pipettes (5-9 MΩ) were pulled from borosilicate glass pipettes (Sutter Instruments) using a vertical 2-stage puller (Narishige PP-830) and filled with an internal solution containing (in mM): K-gluconate 145, HEPES 10, EGTA 1, Mg-ATP 2, Na2-GTP 0.3, and MgCl2 2 (Thermo Fisher). This solution was adjusted to 7.3 pH and 290-300 mOsm, and 0.5% biocytin was added on the day of the experiment.

Patch-clamp electrophysiology was conducted using a Nikon E600 upright microscope and a 60X water immersion objective, and slices were held in a chamber containing recording ACSF with a flow rate of 2-3 mL/min at room temperature. The PL was identified by its anatomical position between the ventral endopiriform cortex, striatum and basolateral amygdala and its higher cell density than nearby regions.

Whole-cell patch-clamp recordings were obtained using a MultiClamp 700A amplifier, Digidata 1550B digitizer, and pClamp 10 software (Axon Instruments, Molecular Devices). Electrical signals were sampled at 20 KHz. During whole-cell current clamp recordings, cells with access resistances greater than 30 MΩ or resting membrane potential drift >5 mV were discarded. 1-second long hyperpolarizing and depolarizing current steps were applied, in increasing intervals of 10 pA up to ±200 pA. Next, voltage clamp recordings were obtained to record sEPSCs (inward currents at -70 mV holding potential) and sIPSCs (outward currents at -40 mV holding potential). The following steps were taken to facilitate biocytin filling: each whole-cell configuration was maintained for at least 10 minutes, after which the pipette was slowly removed from the cell surface over a course of 2-3 minutes. Slices remained in the chamber for another 15 minutes before proceeding to fixation (see Tissue Processing section below).

### Surgeries

All procedures were conducted in accordance with and approved by the Institutional Animal Care and Use Committee at Children’s National Hospital. For all surgeries, C57/Bl6 mice (Jax #000664) (P50-80) were anesthetized with isoflurane and placed into a stereotaxic apparatus (Stoelting co. #51600). Body temperature was maintained with a heating pad during surgery and recovery, and 1-2% isoflurane was delivered continuously through a nose port. Animals were treated with analgesic buprenorphine (0.09 mg/kg body weight of 0.03 mg/mL buprenorphine prepared in sterile saline) prior to surgery, and every 12 hours afterwards as needed. Mice were monitored daily and sacrificed 3-4 weeks post viral injection.

All injection coordinates were measured from Bregma. For all injections, the syringe remained in place for 2-4 min following delivery of virus and then slowly withdrawn. For retrograde tracing of PL inputs, AAV2-retro virus (rAAV1-retro-EF1a-DO_DIO_TdTomato_EGFP-WPRE-pA, Addgene #37120, 1 x 10^13^ GC/mL, (Tervo et al., 2016) was injected bilaterally (50nL at 20 nL/min) into the PL (AP: -0.4, ML: ±2.8, DV: -4.75). For anterograde tracing of input sources, rAAV5-hsyn-mCherry-WPRE (Addgene #114472, Titer: 8 x 10^12^ GC/mL) was injected unilaterally (60-100 nL at 40 nL/min) to the following coordinates: BLA (AP: -1.5, ML: ±2.9, DV: -4.7), COA (AP: -2.4, ±2.9, -5.4), nLOT (AP: -0.6, ML: ±1.9, DV: -5.5), ENT (AP: -4.6, ±3.2, -3.6), AIC (AP: 2.2, ML: ±2.1, DV: - 2.6). For anterograde tracing of PL outputs, AAV1-phSyn1(S)-FLEX-TdTomato-T2A-Syp-EGFP (Addgene #51509, 4 x 10^14^GC/mL) was injected (60nL at 20nL/min) to adult (P54-112) *VGlut1-ires-cre* mice (Jax #037512).

### Tissue processing

Mice were anesthetized with isoflurane and transcardially perfused with 20 mL of 1X phosphate buffered saline (PBS) followed by 20 mL of fixative (4% paraformaldehyde in 1X PBS). Following perfusion, brains were kept overnight in fixative at 4°C with shaking, and cryoprotected in a solution containing 30% sucrose in PBS for 24-48 hours at 4°C with shaking. Next, whole brains were embedded in OCT compound (Fisher Healthcare) and kept at -80°C for at least 24 hours. Sagittal sections were cut at 50µm using a cryostat (Leica CM3050S) and stored in PBS containing 0.02% sodium azide.

For immunolabeling, sections were washed in PBS with 0.2% Triton (PBST) and then incubated for 2 hours at RT in a blocking solution containing PBST with 5% Normal Donkey Serum. After blocking, primary antibodies were added to fresh blocking solution, and sections were incubated overnight at 4°C. The following primary antibodies were used: Rabbit anti-Tbr1 (1:500, EMD Millipore 10554), goat anti-Tdtomato (1:500, Sicgen AB8181), rat anti-GFP (1:1000, Nacalai-Tesque 0404), guinea pig anti-VGlut2 (1:500, Millipore AB2251). Sections were then washed with PBST, 3 times for 5 minutes per wash, and 2 times for 10 minutes per wash. Then, sections were incubated in secondary antibodies in fresh blocking solution, for 3 hours at room temperature. The following secondaries were used: Donkey anti-chicken Alexa-Fluor 647 (1:1000, Jackson Immuno 703-605-155), donkey anti-goat Cy3 (1:1000, Jackson Immuno 705-165-003), donkey anti-rat FITC (1:1000, Jackson Immuno 712-095-150), donkey anti-guinea pig Alexa-Fluor 488 (1:1000, Jackson Immuno 706-545-148). Following secondary incubation, sections were again washed with PBST, 3 times for 5 minutes per wash, and 2 times for 10 minutes per wash. Finally, sections were mounted to slides and coverslipped with Fluoromount-G with DAPI (SouthernBiotech).

For post-hoc biocytin labeling of patch-clamp slices, after recordings sections were fixed in 4% PFA overnight at 4°C, and then washed in PBST. Fixed sections were incubated overnight at 4°C in fresh PBST containing Streptavidin-Cy5 (1:500, Vectorlabs SA-1500-1), and then washed with PBST 3 times for 10 minutes per wash, and 2 times for 30 minutes per wash. Finally, sections were mounted to slides, allowed to dry completely, and coverslipped with Fluoromount-G with DAPI (SouthernBiotech).

### Microscopy

All imaging was performed using a Nikon A1 confocal microscope. Biocytin-filled neurons were imaged using 10x (NA 0.4), 20x (NA 0.75), and 60x (NA 1.4) objectives. Z-stack mosaics (Z-step 1.05 µm) acquired at 20x were used for morphological reconstruction, and 60x Z-stacks (Z-step 0.125 µm) were used for soma reconstruction. For viral-traced tissue, 10x mosaics of whole slides were first acquired with a large pinhole size (45.8µm), followed by 20x (NA 0.75) and 40x (NA 1.3) Z-stacks of regions of interest. Quantification of input neuron sources (Figure 3) was performed by counting labeled neurons within each region using 10x objective in epifluorescence mode. To visualize VGlut2-expressing synaptic terminals within the PL (Figure 4), the 60x objective was used with an additional scan zoom of 2.52X to achieve maximum magnification (0.11 µm per voxel).

### Experimental Design and Statistical Analysis

For patch clamp electrophysiology, n=37 neurons from n=24 mice (12 females/12 males, P59-78) from n=13 different litters were used. Only one recorded neuron per brain section was used, to prevent ambiguity in morphological renderings. Analysis was performed in Clampfit 10. Input resistance was calculated from the linear slope of the voltage responses to the first 3-4 hyperpolarizing current steps. Time constant was calculated from an exponential curve to a 10-pA hyperpolarizing current. Sag ratio was defined as the ratio of the steady state voltage to the peak negative voltage reached during a hyperpolarization step. Action potential threshold, latency, peak, amplitude, spike duration, upslope, and downslope were analyzed from the first elicited action potential (at the rheobase current). Threshold was measured as the first point with dV/dt > 20 mV/msec. latency was defined as the time between the beginning of the current step and threshold. Amplitude was calculated as the difference between peak voltage and threshold. Spike duration was measured as the width of the spike halfway between threshold and peak. Upslope/downslope ratio was defined as the ratio between the maximum upslope and downslope, which were calculated as the largest and smallest values (respectively) of dV/dt during the action potential. Burst index was measured using the voltage response to a depolarizing current 50 pA greater than rheobase for each neuron. It was calculated as 1-F0/F1, where F0 is the frequency of the beginning of the spike train, and F1 is the frequency at the end of the spike train.

Following analysis, agglomerative hierarchical clustering (with Ward linkage and Euclidean distance) was performed using the matplotlib package (python) (Figure 1). The following input variables were used for clustering: resting membrane potential, input resistance, time constant, and sag ratio, as well as action potential latency, rheobase, threshold, amplitude, duration, burst index, and upslope/downslope ratio. Capacitance, action potential peak voltage, upslope, and downslope were excluded from clustering. Clustering results were used to compare each individual physiological and morphological parameter between clusters. For these comparisons, unpaired t-tests were used, and the alpha cutoff was adjusted to 0.01 to correct for multiple comparisons.

**Figure 1.**
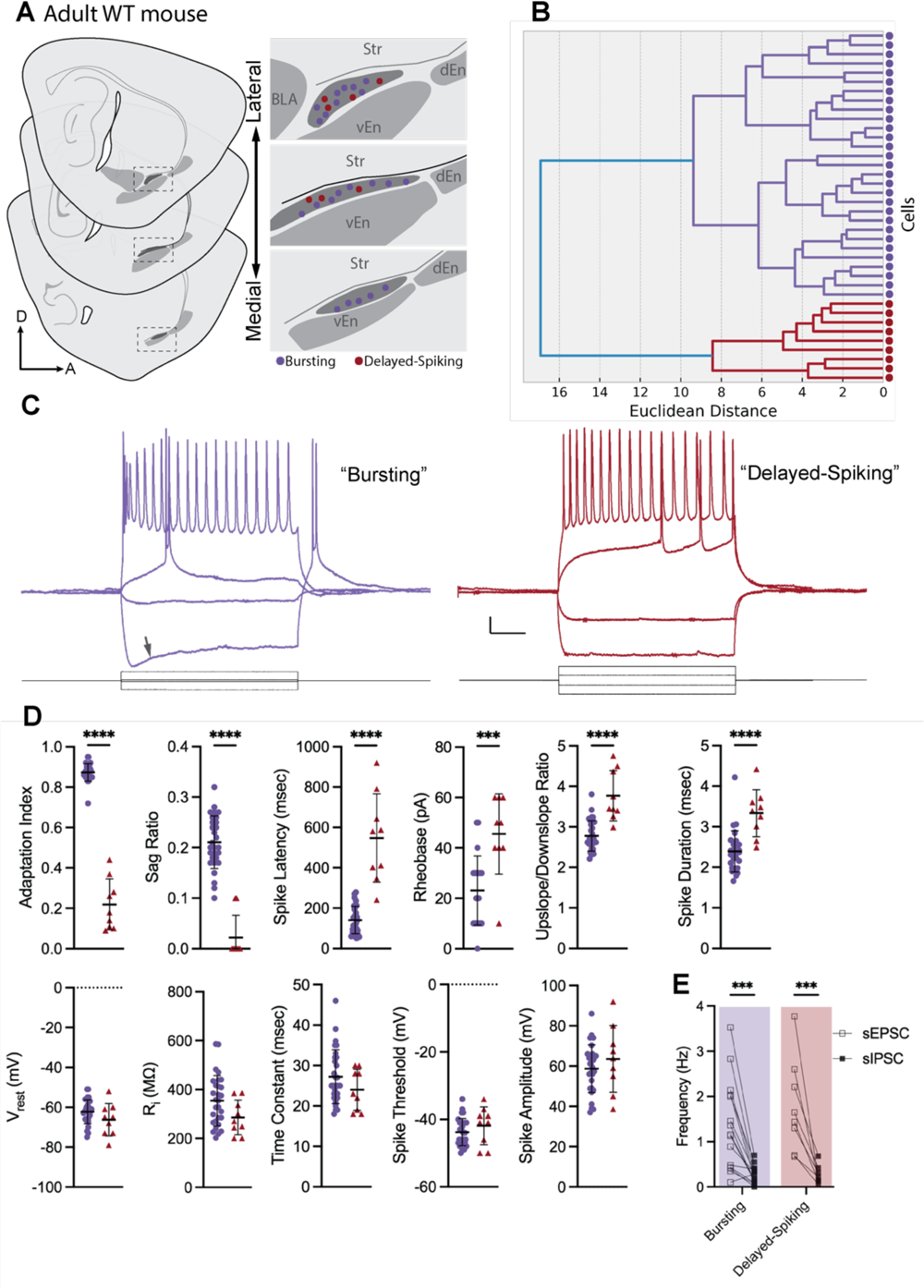
Electrophysiological subtypes in the adult mouse paralaminar amygdala. (A) Patch-clamp electrophysiology was performed in sagittal sections from adult (P59-78) WT mice targeting neurons across the PL. (B) An agglomerative hierarchical clustering dendrogram depicts two clusters of cells (purple and red) separated by a Euclidean distance of 17. (C) Neurons within each cluster exhibited stereotyped action potential patterns in response to 1-second current injections. Arrow indicates anomalous rectification exhibited by bursting neurons during hyperpolarization (see main text). (D) T-test comparisons between subtypes of each individual electrical parameter used for clustering, with alpha-cutoff adjusted to 0.01 to correct for multiple comparisons. (E) Frequencies of spontaneous excitatory (sEPSC) and inhibitory (sIPSC) postsynaptic currents. Scale bars in (C): 10mV (vertical) and 200msec (horizontal). *** p < 0.0001, *** p < 0.001.

Morphological analysis (Figure 2) was performed in Imaris 10. Each neuron was reconstructed using the ‘Filaments’ tool in the 3D view, and the soma was rendered using the ‘Surfaces’ tool. Sholl analysis was performed using concentric radii of 20µm each. Branch frequency was calculated by dividing the number of branch points by the total filament length for each neuron.

**Figure 2.**
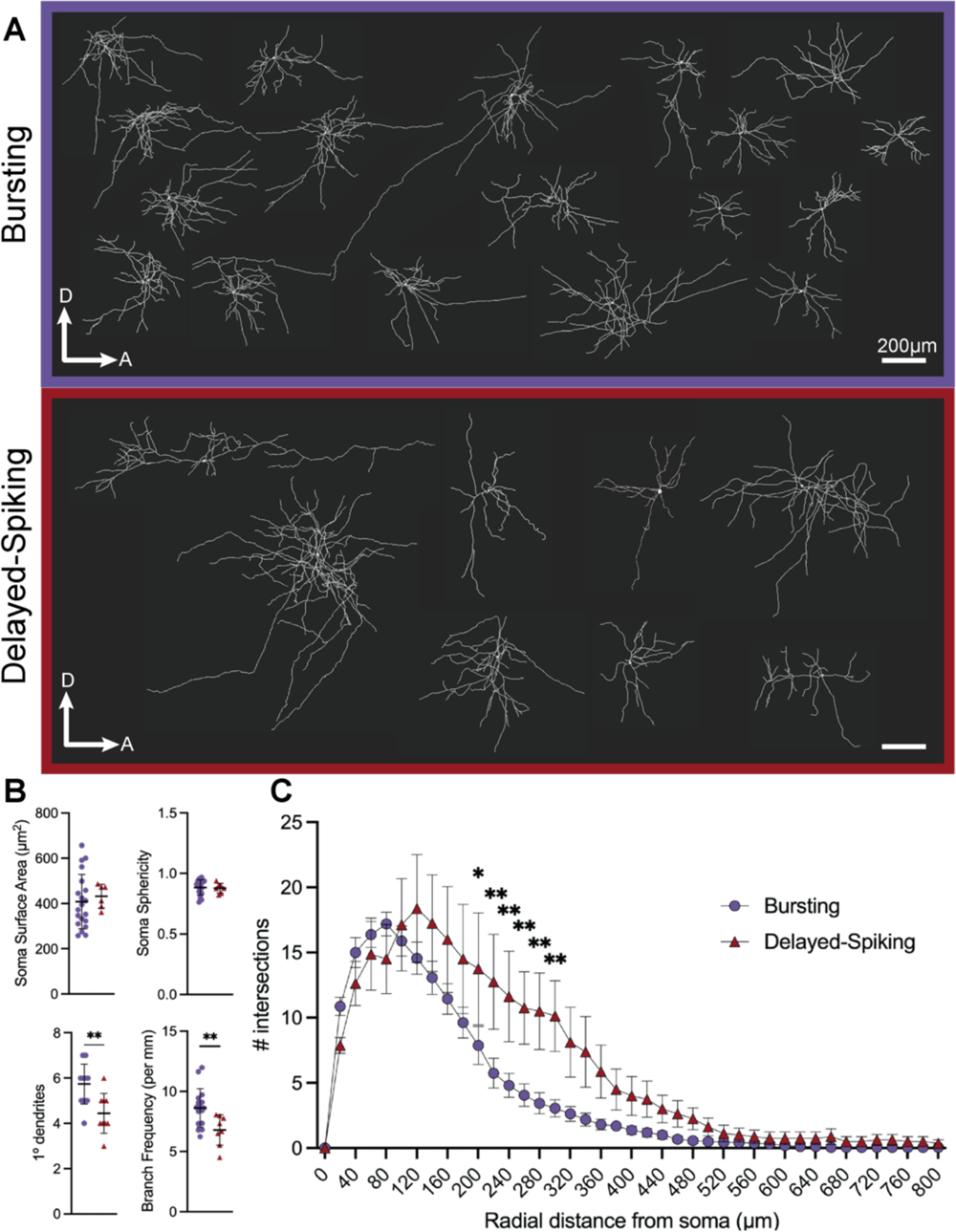
Bursting and delayed-spiking PL neurons possess distinct morphologies. Following patch-clamp recordings, post-hoc biocytin staining and morphological reconstructions (A) were performed on bursting (purple) and delayed-spiking (red) neurons. (B) Comparisons between subtypes of soma surface area and sphericity, number of primary dendrites (dendrites directly emanating from the soma), branch frequency (number of branches per filament distance) data are Mean ± SD. (C) Number of Sholl intersections in each subtype at 40μm radial intervals (Mean ± SD). ** p <0.01.

Evaluation of sex differences in morpho-electric criteria (Extended data Figure 1-1) was performed using z-scores of each electrophysiological and morphological variable. Z-scores were obtained using the fit_transform() function in the matplotlib package (python). This transformed data was analyzed using multiple t-tests, correcting for multiple comparisons using the two-step false discovery rate (FDR) approach, with FDR equal to 1%.

For retrograde tracing of PL inputs (Figure 3), n=6 mice (2 males, 4 females) with successful PL targeting were analyzed. To control for variable transduction across mice, labeled neurons in each input region were not directly compared between mice, but instead compared as a percentage of total input neurons within each mouse. For anterograde tracing of input sources projecting to the PL (Figure 4), n = 3 mice per input source were analyzed (2 females, 1 male for COA, nLOT, ENT, AIC, and 2 males 1 female for BLA). Analysis was done by confocal visualization of labeled fibers within the PL colocalized with VGlut2 expression. For anterograde tracing of output targets emanating from the PL (Figures 5-8), n = 4 mice (3 males, 1 female) with successful targeting were analyzed. Analysis was done using Nikon NIS-Elements software using a 10x large image stitch of an entire slide. For each output target region, multiple ROIs were drawn across sections, and the average intensity of the FITC channel (Syp-EGFP) was measured. The vEN and dEN regions were excluded from analysis due to viral leakage from the adjacent PL. To control for variable transduction and histology, intensity values within each mouse were normalized to percentages, using 0% as the lowest observed intensity and 100% as the highest observed intensity (Graphpad Prism 10).

**Figure 3.**
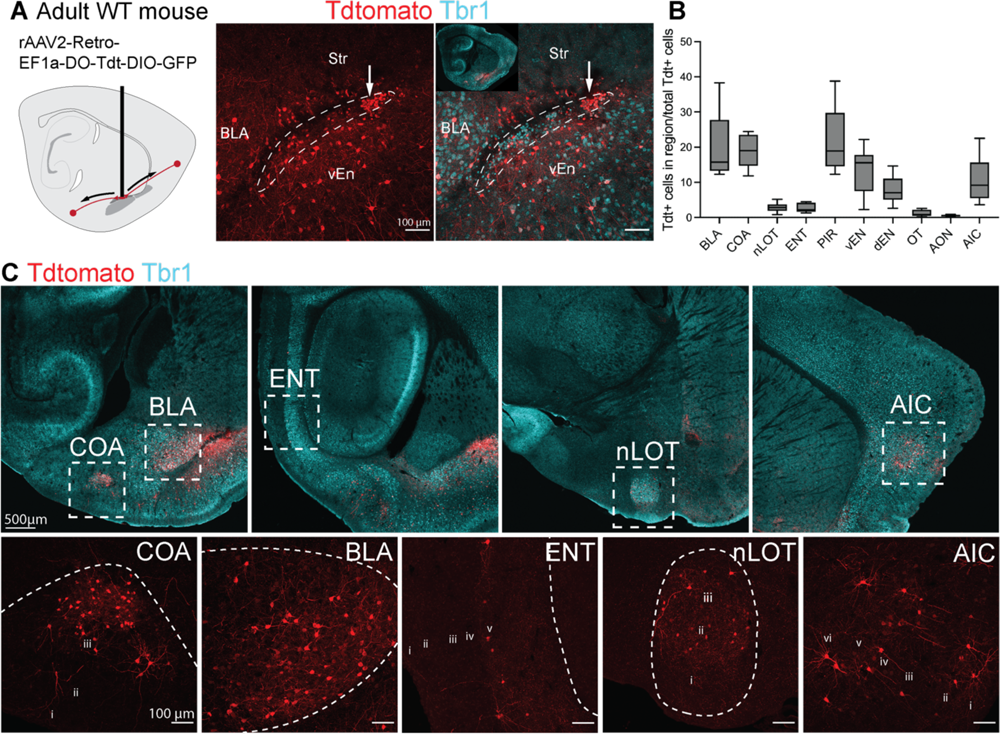
Retrograde viral labeling reveals putative regions of input to the PL. (A) rAAV2-Retro-EF1a-DO-Tdt-DIO-GFP was injected to the PL of adult (P50-64) WT mice. White arrow indicates injection site. (B) Quantification of tdTomato+ labeled neurons within each brain region as a percentage of total labeled neurons for each experiment. (C) Input neurons located within the cortical and basolateral amygdala, entorhinal cortex, nucleus of the lateral olfactory tract, and agranular insular cortex.

**Figure 4.**
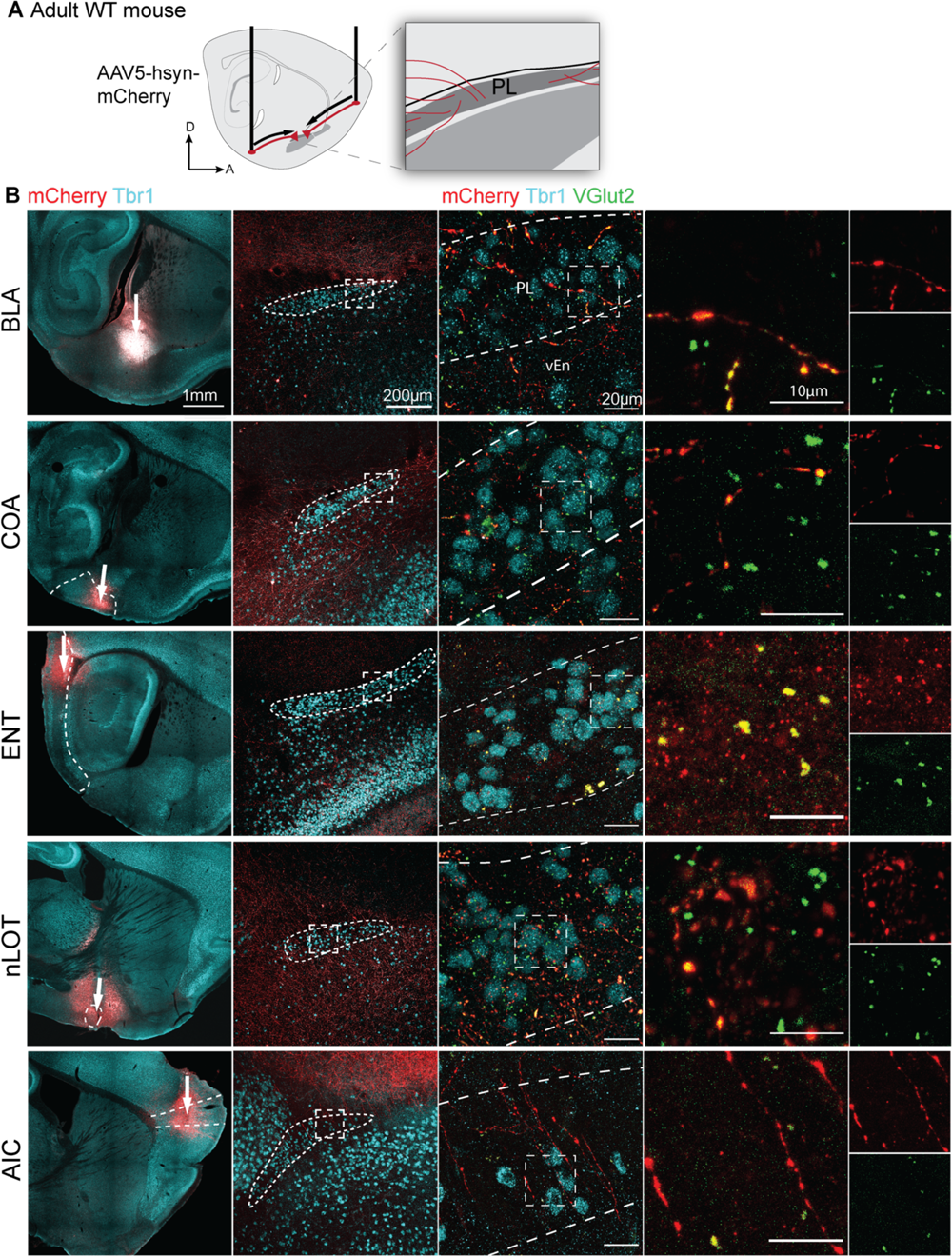
Anterograde viral labeling of inputs to the PL. (A) Major regions of input uncovered in Figure 3 were infected with AAV5-hsyn-mCherry in adult (P54-68) WT mice. (B) Injections to either the BLA, COA, ENT, nLOT, or AIC (left, white arrows indicate injection sites), and labeled fibers (mCherry) within the PL following each injection (center). Labeled fibers co-localized with VGlut2+ putative excitatory presynaptic terminals (far right).

**Figure 5:**
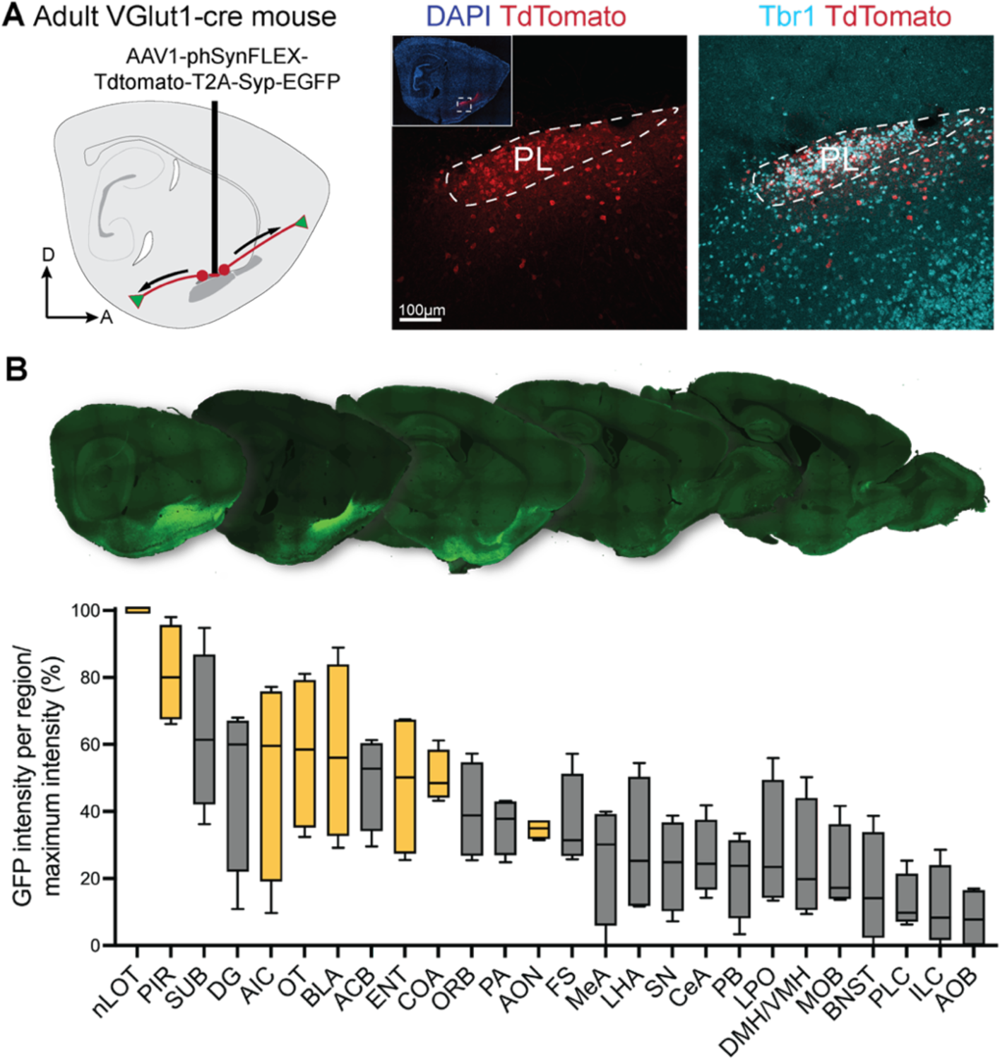
Anterograde viral tracing revealing the output projections from the adult mouse PL. (A) AAV1-phSyn1(S)-FLEX-tdtTomato-T2A-SypEGFP-WPRE was injected to the PL of adult (P54-112) *VGlut1-cre* mice. (B) Example sagittal sections and quantified mean intensity of Syp-EGFP labeling within brain regions. EGFP intensity was normalized in each mouse, with 100% and 0% corresponding to the highest and lowest observed intensities respectively. Yellow boxes indicate regions with reciprocal projections to the PL.

**Figure 6.**
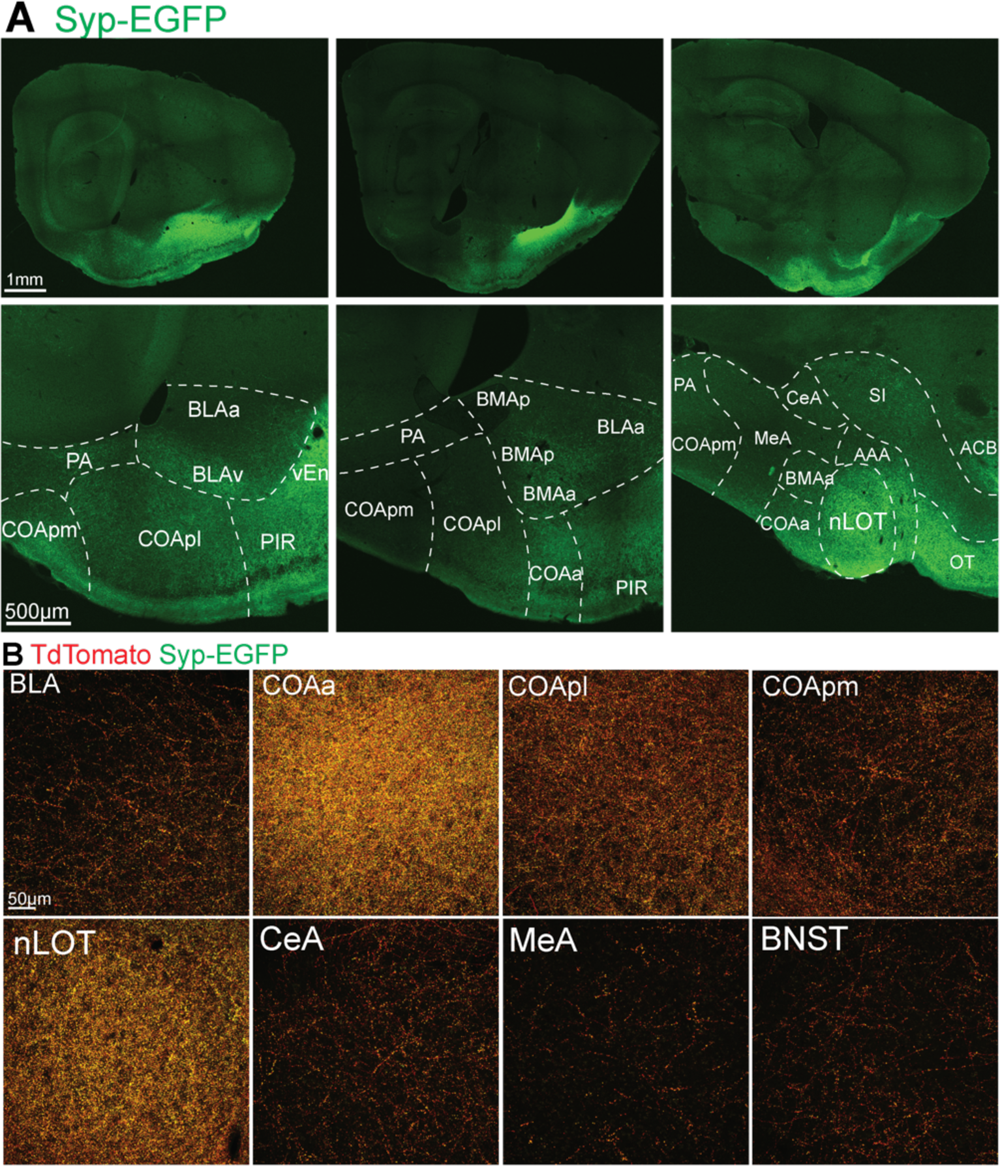
Anterograde viral labeling of PL outputs in the amygdala. (A) Example Syp-EGFP labeling across the amygdala. (B) High magnification images of PL output fibers (tdTomato) and synaptic terminals (EGFP) within amygdala subnuclei.

**Figure 7:**
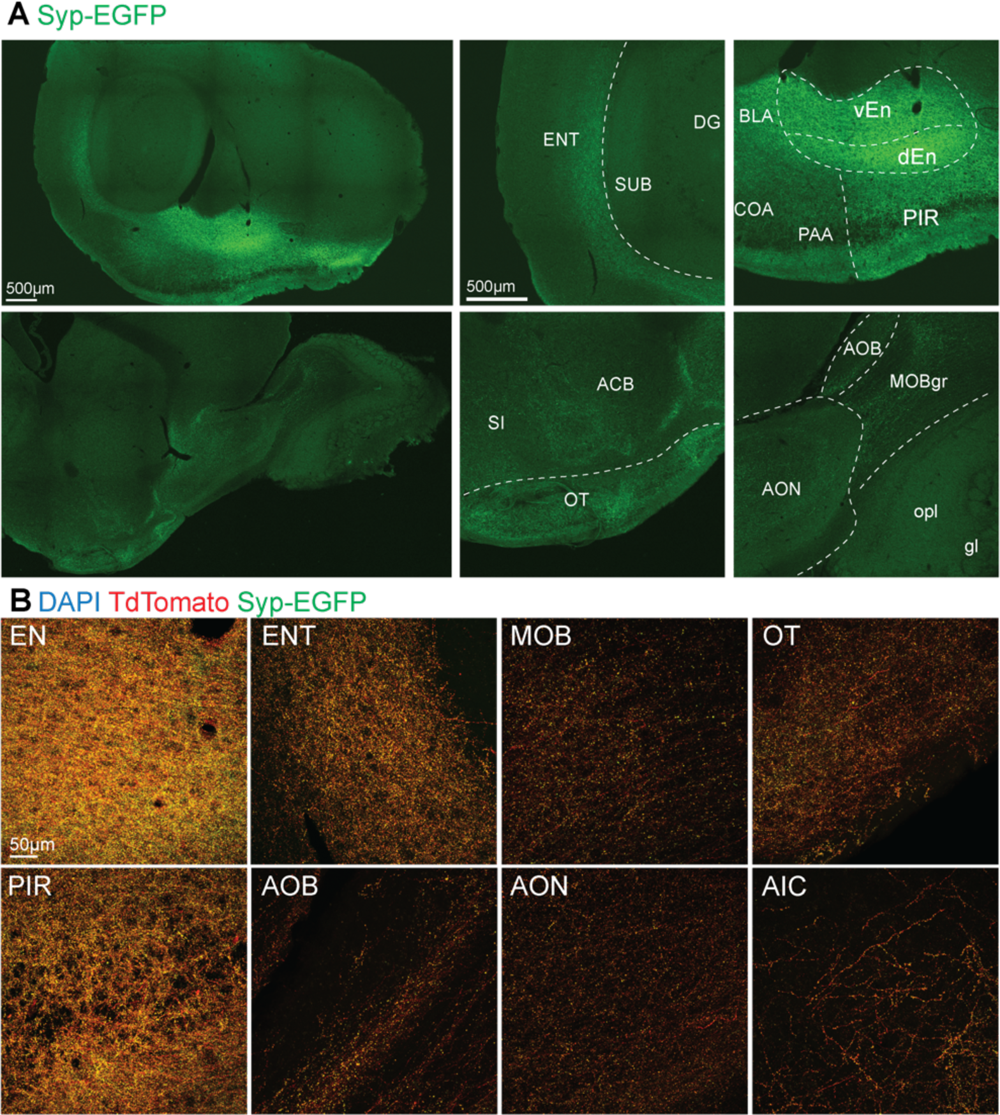
Anterograde viral labeling of PL outputs in the olfactory network. (A) Example Syp-EGFP labeling across olfactory network regions. (B) High magnification images of PL output fibers (tdTomato) and synaptic terminals (EGFP) within olfactory network regions.

**Figure 8:**
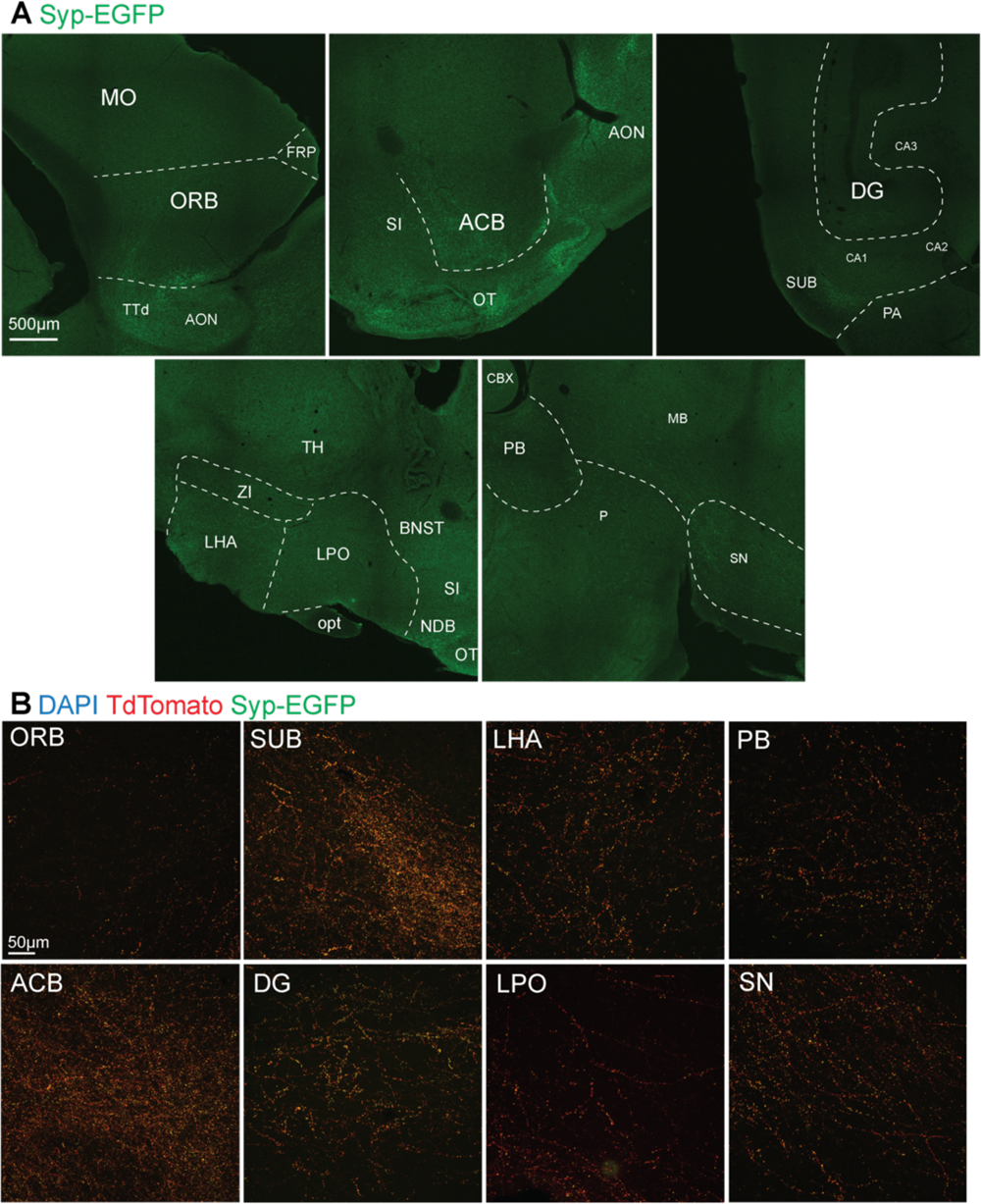
Anterograde viral labeling of PL outputs in frontal, striatal, hippocampal, hypothalamic, and brainstem regions. (A) Example Syp-EGFP labeling. (B) High magnification images of PL output fibers (tdTomato) and synaptic terminals (EGFP) within frontal, striatal, hippocampal, hypothalamic, and brainstem regions.

**Table 1.** List of Abbreviations.

|  |  |  |  |
| --- | --- | --- | --- |
| AAA | Anterior Amygdala Area | LPO | Lateral Preoptic Area |
| ACB | Nucleus Accumbens | MB | Midbrain |
| AIC | Agranular Insular Cortex | MeA | Medial Amygdala |
| AOB | Accessory Olfactory Bulb | MO | Somatomotor Cortex |
| AON | Anterior Olfactory Nucleus | MOB | Main Olfactory Bulb |
| BLA | Basolateral Complex of the Amygdala | gr | granule layer |
| BNST | Bed Nucleus of the Stria Terminalis | opl | outer plexiform layer |
| CBX | Cerebellar Cortex | gl | glomerular layer |
| CeA | Central Amygdala | nLOT | Nucleus of the Lateral Olfactory Tract |
| COA | Cortical Amygdala | opt | Optic Tract |
| COAa | Anterior Cortical Amygdala | ORB | Orbitofrontal Cortex |
| COApl | Posterolateral Cortical Amygdala | OT | Olfactory Tubercle |
| COApm | Posteromedial Cortical Amygdala | P | Pons |
| DG | Dentate Gyrus of the Hippocampus | PA | Posterior Amygdala |
| DMH | Dorsomedial Hypothalamus | PB | Parabrachial Nucleus of the Pons |
| EN | Endopiriform cortex/nucleus | PIR | Piriform Cortex |
| vEN | Ventral Endopiriform | PL | Paralaminar Nucleus of the Amygdala |
| dEN | Dorsal Endopiriform | PLC | Prelimbic Cortex |
| ENT | Entorhinal Cortex | SI | Substantia Innominata |
| FRP | Frontal Pole | SN | Substantia Nigra |
| FS | Fundus of Striatum | Str | Striatum |
| ILC | Infralimbic Cortex | SUB | Subiculum |
| ITC | Intercalated Cells of the Amygdala | TT | Tenia Tecta |
| LHA | Lateral Hypothalamic Area | VMH | Ventromedial Hypothalamus |

## Results

### Electrophysiological subtypes of adult PL neurons

The adult mouse PL contains excitatory neurons, but their diversity is unstudied. To investigate the electrical properties of these cells, we performed patch-clamp electrophysiological recordings from individual PL neurons in adult (P59-78) C57/Bl6 mice (Fig 1). We measured voltage responses to 1-second hyperpolarizing and depolarizing current steps of increasing magnitude in n = 37 neurons (from n = 24 mice) sampled from multiple levels across the PL (Fig 1A). We performed unsupervised agglomerative hierarchical clustering using 11 intrinsic electrical parameters (see methods) to generate an unbiased representation of physiological diversity among the recorded cells. The resulting dendrogram segregated cells into two main clusters (Fig 1B). Further investigation revealed that this segregation was due to the different action potential firing patterns between the clustered cells (Fig 1C). All neurons within the larger cluster (n = 28 cells) exhibited initial bursts of 2-3 action potentials during depolarization, and anomalous rectification during hyperpolarization. In the smaller cluster, all neurons (n = 9) exhibited spike latencies of several hundred milliseconds during depolarization, and no rectification during hyperpolarization. Based on these stereotyped patterns, we labelled these clusters “bursting” and “delayed-spiking” neurons.

To better understand the differences between bursting and delayed-spiking PL neurons, we performed unpaired t-tests of each physiological parameter used for hierarchical clustering individually (Fig 1D). There were no significant differences in passive membrane properties (RMP t(36)=1.59, p=0.1209; input resistance t(36)=1.85, p=0.0729; time constant t(36)=1.34, p=0.1893; capacitance t(36)=0.948, p=0.3495), action potential threshold (t(36)=1.12, p=0.2698), or amplitude (t(36)=0.962, p=0.3425). There were statistically significant differences in adaptation index (t(36)=24.2, p<0.0001), spike latency (t(36)=8.92, p<0.0001), and sag ratio (t(36)=9.76, p<0.0001) between subtypes, which reflected the distinct properties we observed in action potential patterns (Fig 1C). This analysis further revealed that bursting neurons displayed significantly lower rheobase (t(36)=4.15, p=0.0002), spike duration (t(36)=4.69, p<0.0001), and upslope/downslope ratio (t(36)=5.87, p<0.0001). Together, these properties define two distinct electrophysiological subtypes of neuron in the adult mouse PL.

In a subset of neurons, we measured spontaneous synaptic input activity (n = 23) (Fig 1E). There was no significant difference between excitatory (sEPSC) and inhibitory (sIPSC) postsynaptic current frequencies in either bursting or delayed-spiking neurons (sEPSC t(42)=1.379, p=0.1751; sIPSC t(42)=0.04251, p=0.9663, mixed effects analysis with post-hoc Fisher’s LSD). Interestingly, however, both subtypes received significantly greater frequency of excitatory input than inhibitory input (bursting t(42)=4.066, p=0.0002, delayed spiking t(42)=4.215, p=0.0001, mixed effects analysis with post-hoc Fisher’s LSD). Together these data suggest that both bursting and delayed-spiking neurons receive synaptic inputs.

Based on our prior work showing that PL neurons develop along a delayed trajectory, the possibility exists that the electrophysiological subtypes we observe are due to different stages of maturation of PL neurons. Our prior work indicates that by molecular criteria, ∼90% of neurons in the adult (P60) mouse PL are mature (Dcx-, NeuN+). Further, in our previous recordings of at juvenile ages (P21-28), we observed that electrophysiological maturation of PL neurons is defined by the combination of decreasing input resistance, hyperpolarizing threshold, and reduction in spike duration (Alderman et al., 2023). Compared to those juvenile PL neurons we previously recorded, the adult PL neurons we recorded here exhibit lower input resistance (338 mV in adult, 813 MΩ in juvenile PL), hyperpolarized threshold (-43 mV in adult, -40 mV in juvenile PL), and shorter spike duration (2.61 msec in adult, 3.61 msec in juvenile PL). To further investigate possible differences in maturity between adult subtypes, we plotted this combination of features (Extended data Fig 1-1A). This revealed that in contrast to the juvenile PL neurons in our prior studies, adult PL neuron physiological properties do not fall on a maturational continuum. Further, across the adult age range we recorded (P59-78), none of the criteria that define juvenile PL neuron maturational stage (input resistance, threshold, spike duration) were correlated with animal age (input resistance r=-0.2, threshold r=-0.02, spike duration r=-0.07, Pearson’s r, Extended data Fig 1-1B). This data indicates that electrophysiological differences in adult PL subtypes are not based on different maturational stages.

### Adult PL neuron electrophysiological subtypes have distinct morphologies

We next examined whether there were morphological differences between bursting and delayed-spiking PL neurons (Fig 2). During patch clamp recordings, n = 19/29 bursting neurons and n = 8/9 delayed-spiking neurons were filled with biocytin, and we post-hoc reconstructed their morphologies in 275 µm thick sections using Imaris. The anatomical location and orientation of each reconstructed cell was also determined. The delayed-spiking neurons we recorded were primarily located in lateral PL sections with none observed in the medial PL, whereas the bursting neurons we recorded were distributed throughout the PL (Fig 1A). In each subtype, the reconstructed neurons exhibited substantial variety in morphology (Fig 2A), similar to our previous findings in the juvenile PL where morphological complexity varied independently of physiological maturity (Alderman et al., 2023).

To determine whether bursting and delayed-spiking neurons exhibit differences in soma size, shape, or dendritic architecture, we analyzed these properties in Imaris. The somas of bursting and delayed-spiking neurons were of similar size (t(24)=0.459, p=0.6504) and sphericity (t(24)=0.245, p=0.8082), but bursting neurons had a greater number of primary dendrites (t(24)=3.46, p=0.0022) as well as a higher branch frequency (t(24)=2.83, p=0.0098, unpaired t-tests) (Fig 2B). Sholl analysis revealed that delayed-spiking neurons exhibited a greater number of sholl intersections from 200-300µm radii (200µm p=0.0413, 220µm p=0.0038, 240µm p=0.0059, 260µm p=0.0077, 280µm p=0.0033, 300µm p=0.0029, mixed-effects ANOVA with Sidak correction) (Fig 2C). Thus, while adult PL neurons exhibit diverse morphologies, bursting and delayed-spiking subtypes exhibit significant differences in dendritic complexity.

The amygdala is a sexually dimorphic structure. To investigate whether neurons in the adult PL exhibit sex differences, we compared the physiological and morphological features collected from male (n = 12) and female (n = 12) mice. Across all features, we observed no significant differences between sexes (multiple unpaired t-tests with FDR of 1%, Extended data figure 1-1C).

From these combined observations, we conclude that in contrast to prior molecular data suggesting homogenous excitatory neurons, the adult PL instead contains two neuronal subtypes defined by distinct action potential firing patterns and morphologies. In both subtypes, we recorded a similar frequency of spontaneous synaptic input; we next investigated the sources of this input to the PL.

### The mouse PL receives major input from the main olfactory network and amygdala basolateral complex

To uncover the sources of neuronal input to the mouse PL we performed unilateral injections in the PL of adult WT mice (P50-64, n = 6) with the retrograde virus *rAAV2-retro-EF1a-DO-TdTomato-DIO-EGFP* (Addgene #37120) (Fig 3A). Following 2-3 weeks of incubation, we observed tdTomato+ neurons in multiple regions within and outside the amygdala (Fig 3B, C). We quantified the relative locations of these cells within each injected brain by normalizing the number of tdTomato+ cells in each region to the total number infected per experiment (Fig 3B). Across all experiments, labeled neurons were observed in multiple nuclei of the main olfactory network (anterior olfactory nucleus (AON), piriform cortex (PIR), olfactory tubercle (OT), cortical amygdala (COA), nucleus of the lateral olfactory tract (nLOT), entorhinal cortex (ENT), and agranular insular cortex (AIC)), as well as in the amygdala basolateral complex (BLA) (Fig 3B).

The relatively small size of the PL makes it challenging to target specifically without viral spread to the nearby ITCs, external capsule, and vEN. We qualitatively assessed the extent of this spread in our injections and found that similar input regions were identified independently of which neighboring regions included viral spread (Extended data Figure 3-1).

Although our input observations were consistent across injections, many of the regions we observed also project to the nearby piriform and endopiriform cortex (Majak & Moryś, 2007; Neville & Haberly, 2004). Therefore, we next investigated whether these putative input sources send projections specifically into the PL. To accomplish this, we selected a subset of regions that contained a substantial proportion of putative input neurons (> 1% of total tdT+ cells). In these selected input regions (BLA, COA, nLOT, ENT, and AIC) we performed unilateral injections in adult WT mice (P57-67) with the anterograde virus *rAAV5-hsyn-mCherry*-*WPRE* (Addgene #114472). Each input region was targeted in an individual experiment (Extended data figure 4-1). Following 3 weeks of incubation, brains were sectioned at 50 µm and stained with Tbr1 to label the PL, mCherry to reveal infected cells and projections, and VGlut2 to label excitatory synaptic terminals (Figure 4). We observed mCherry+ fibers emanating from each input source into the PL. In all experiments, these fibers colocalized with the glutamatergic presynaptic marker VGlut2. Together, these findings indicate that the PL receives excitatory neuronal input from the basolateral complex, cortical amygdala, nucleus of the lateral olfactory tract, entorhinal cortex, and agranular insular cortex, suggesting a role in integrating higher-order olfactory and valence information.

### The adult mouse PL projects reciprocally to its input sources and to the hippocampus, striatum, hypothalamus *<u>and brainstem</u>*

We next investigated which brain regions are targeted by PL output projections. The mouse PL expresses the glutamatergic marker Slc17a7 (VGlut1), distinguishing it from the neighboring inhibitory ITCs and ventral striatum (Alderman et al., 2023). We performed injections in the PL of adult (P54-112) *VGlut1-ires-cre* transgenic mice (Jax #037512, n = 4) with the anterograde virus *AAV1-phSyn-FLEX-Tdtomato-T2A-Syp-EGFP* (Addgene #51509), which, in the presence of cre recombinase, labels infected neurons with tdTomato and synaptic terminals with EGFP (Fig 5A) (Oh et al., 2014). As the nearby basolateral complex and piriform/endopiriform cortex also contain VGlut1-expressing cells, we qualitatively scored their degree of infection (Extended data figure 5-1).

Following 3 weeks of incubation, we performed sagittal sectioning of whole brains, and quantified the intensity of GFP (putative synaptic terminals) in each observed brain region (Fig 5B). Because of the leakage of virus to the endopiriform cortex, it was excluded from quantification. To evaluate whether the GFP signal intensity observed corresponded to synaptic terminals labeled by viral infection, we verified the colocalization of GFP with tdTomato+ labeled fibers in every observed target region (Figs 6, 7, 8). Across all regions, synaptic terminals (GFP) colocalized with fibers (tdTomato).

Strong GFP signal was present in all PL input regions (Fig 5B) observed in our retrograde tracing experiments (Figure 4). The strongest signal across mice was present in the nLOT, and further examination of the amygdala revealed dense terminals in the basolateral and cortical nuclei (Figure 6). In the COA, terminals were densest in the anterior subregion. Investigation of the olfactory network confirmed wide targeting of all structures by labeled terminals, including input sources (ENT, EN, PIR, OT, AON, AIC), as well as the main and accessory olfactory bulbs (MOB, AOB) (Figure 7). In addition to the amygdala and olfactory network, the hippocampal subiculum (SUB) and dentate gyrus (SG) exhibited strong signal, as well as the nucleus accumbens (ACB) (Figs 5B, 8). Finally, labeled fibers traveled via the hypothalamus and into the brainstem (Figure 8), with putative synaptic terminals present in the lateral and medial hypothalamus, substantia nigra, and parabrachial nucleus (Fig 5B).

Together, these results indicate that the outputs of the PL emanate primarily to olfactory, amygdala, hippocampal, and striatal structures, with additional projections to hypothalamus and brainstem. This suggests that the PL may influence olfactory, affective, and contextual learning, as well as autonomic and visceral states.

## Discussion

Using a combination of electrophysiological, morphological reconstruction and viral based circuit mapping, we identified bursting and delayed-spiking neuronal subtypes in the adult mouse PL, as well as mapped its input and output projections. The PL is reciprocally connected with olfactory regions, with additional outputs primarily to valence processing and memory structures. Our findings provide critical novel understanding of this understudied population, and are essential to elucidate the role of this novel population in complex brain and behavior changes that unfold during the transition from adolescence to adulthood, and their role in the adult brain.

Action potential patterns are a common distinguishing feature of neuronal subtypes. PL subtypes are differentiated by the presence or absence of bursting, a property which frequently defines divergent circuit roles in both vertebrate and invertebrate organisms (Krahe & Gabbiani, 2004; Lisman, 1997; Zeldenrust et al., 2018). One mechanism for bursting is the backpropagation of an initial spike into the proximal dendrites, activating dendritic conductances that cause an afterdepolarization (ADP); in the presence of a low-threshold spike, this ADP crosses threshold and results in a burst (Häusser et al., 2000; Izhikevich, 2007). The greater dendritic complexity in the bursting subgroup lends credence to this possibility. Further, in physiological conditions, bursting may reflect the spatial summation of multiple inputs to different dendrites (Poirazi & Mel, 2001; Zeldenrust et al., 2018). One consequence of a burst is synaptic release potentiation at axon terminals, thereby increasing signaling to postsynaptic targets (Swadlow & Gusev, 2001). In the PL, therefore, the bursting subtype may exhibit stronger transmission to downstream neurons than the delayed spiking subtype. Meanwhile, delayed-spiking neurons require larger and longer current injections to reach threshold, indicating that this subtype may be less excitable *in vivo*. Together, this suggests that PL neuronal subtypes may constitute distinct coding elements as a direct consequence of their electrophysiological differences.

Despite the key dendritic differences in bursting and delayed-spiking neurons, both subtypes exhibit variable morphologies. In prior work in the juvenile PL, we observed that some neurons with immature physiological and molecular features possess surprisingly complex morphologies. Thus, dendritic complexity appears only partly influenced by physiological subtype and cell maturity. However, two likely technical sources of the morphological variance we observe are the uneven biocytin filling across patched neurons, and the inconsistent transection of neurites during slice preparation. Both reduce the recoverable fraction of the total dendritic tree. Thus, we focused on metrics which are robust to these confounds. Soma size and shape, and primary dendrite number do not depend on distal neurites which may be lost. Similarly, branch frequency (branches per mm), which displayed relatively low variability in our dataset, is robust to uneven filling because both branch number and total dendritic length are affected. Uncovering the factors that might influence PL neuron morphology, such as different input and output connectivity or hormonal influences, and how they correlate with subtypes, will be an interesting future direction.

Interestingly, the finding of two excitatory neuron subtypes within the adult PL is similar to the heterogeneity of neuronal types in other late-maturing regions. In the late-maturing piriform cortex, two types of principal neurons exist, defined by the presence or absence of bursting as well as morphological correlates (Suzuki & Bekkers, 2006). Further, these subtypes are differentially connected to olfactory and cortical afferents (Suzuki & Bekkers, 2011). However, whether late-maturing piriform neurons comprise both subtypes is unknown. In the neurogenic dentate gyrus of the rodent, newly born neurons mature into granule cells (GCs) with some distinctions from typically-developed GCs (Cole et al., 2020). A lesser studied dentate population are semilunar granule cells, which exhibit earlier birth and different morpho-electric properties than GCs (Gupta et al., 2020; Save et al., 2019). In addition, although they are not late-maturing, the excitatory principal neurons in the basolateral complex have also been divided into two groups, which exhibit bursting and delayed-spiking features similar to the PL (Pare et al., 1995; Rainnie et al., 1993; Washburn & Moises, 1992). Further, these subgroups are differentially active *in vivo* (Paré & Gaudreau, 1996). This indicates that the subtypes of the adult mouse PL recapitulate patterns of heterogeneity present in other brain regions, and that bursting and delayed-spiking PL neurons likely exhibit differences in circuitry and/or functional recruitment.

Input and output tracing results reveal that the adult mouse PL is reciprocally connected to the main olfactory network and basolateral complex. The main olfactory network is defined by the receipt of direct input from the olfactory bulb, and all regions were revealed as both PL input sources and output projections. The olfactory bulb itself was not a source of direct input to the PL but it was a target of PL output projections, which were localized to the granule layer. Notably, the accessory olfactory network (AOB, MeA, BNST) was not a source of PL input, and received comparatively sparse PL output projections, suggesting that the PL may not be involved in innate behavioral responses to pheromones and odorants. Finally, bidirectional input-output connectivity to the basolateral complex, which encodes innate and learned valence of polysensory stimuli, may confer the attractiveness or repulsiveness of olfactory cues. Thus, the PL likely receives second-order sensory information from olfactory network regions, likely integrating this with valence processing (via BLA connections), and feeds this information back to the olfactory network.

Beyond its reciprocal connectivity to olfactory regions, output projections from the PL further highlight its putative processing of valence (via basal ganglia outputs), a possible role in memory (via hippocampal outputs), and influence over visceral and autonomic states (via pontine and hypothalamic outputs). A key question is whether these connections are overlapping or exclusive in PL subpopulations. Many PL outputs are similar to those of the piriform cortex, which projects to the olfactory and prefrontal cortex, cortical amygdala, and entorhinal cortex, as well as the striatum and hypothalamus (Bekkers & Suzuki, 2013; Johnson et al., 2000; Neville & Haberly, 2004; Price et al., 1991). These output connections appear spatially and functionally organized (Y. Chen et al., 2022), contrasting the long-held belief that they are stochastic (Miyamichi et al., 2011; Schaffer et al., 2018). One possibility is that the PL is similarly organized into distinct circuit motifs, which further extend to its additional outputs. This would place the PL as a hub which associates specific olfactory cues with valence information, possibly supporting specific behavioral response, such as approach/avoidance, autonomic arousal, and memory encoding.

In primates, tracing studies report input fibers emanating to the PL from the ventral hippocampus and lateral amygdala (Fudge et al., 2012; Pitkänen & Amaral, 1998), and output projections from the PL that target the deep amygdala as well as the ventral striatum (Fudge et al., 2002; Fudge & Tucker, 2009). Our connectivity findings in the mouse are similar to these, with the exception of the ventral hippocampus which was not an input to the PL in mouse. However, the dense connectivity of the olfactory cortex to the mouse PL is a striking difference. Due to the lack of detailed tracing studies, it is unknown whether the primate PL displays connections to sensory cortices.

In humans, alterations in the number or molecular signatures of PL neurons are seen in multiple neuropsychiatric disorders. In Autism Spectrum Disorder, PL neurons exhibit alterations in transcripts relevant to maturation, and dysfunctional PL development correlates with a lack of amygdala growth into adulthood (Avino et al., 2018). The adult PL is depleted in military veterans with Post-Traumatic Stress Disorder as well as adults with temporal lobe epilepsy (Ballerini et al., 2022; Morey et al., 2020). Therefore, the PL neuron subtypes and circuitry that we observe here may be sensitive to genetic and environmental insults, and they may confer key circuitry affected by neurological and psychiatric disorders. Further studies in humans, and relevant animal models such as the mouse, will be critical to understand the relevance of the PL to various neuropsychiatric conditions.

In summary, in this study, we reveal that the adult mouse PL contains two neuronal subtypes, is reciprocally connected to the olfactory network, and projects widely to striatal, hippocampal, hypothalamic, and brainstem targets. These findings highlight the significance of the PL across species as a late-maturing amygdala subregion that may serve as a key nexus supporting the behavioral transition to adulthood. Given that synaptic input, mature intrinsic identities, and behavioral responsivity of the PL emerges between juvenile and adult ages, an important next step is to track these changes during adolescence, as well as determine the interrelationships between the development of inputs, intrinsic properties, and behavioral learning.

## Acknowledgments

D.S., P.J.A, S.F.S, S.V, and J.G.C acknowledge the National Institute of Mental Health (NIMH) (F30MH133305, R21MH125367, and R01MH128745). We thank the Corbin, Sorrells, and Vicini lab members for feedback throughout the course of the project.

---

**Extended Data Figure 1-1.**
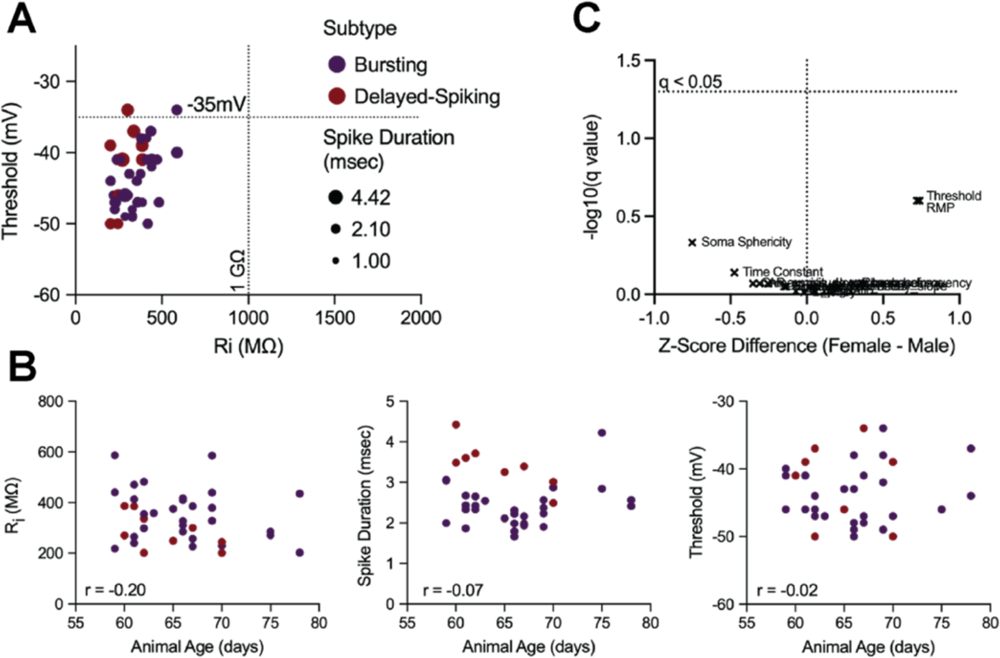
Maturational stages and sex differences in morpho-electric properties of adult (P59-78) mouse PL neurons. (A) Threshold and spike duration plotted against input resistance. Dotted lines represent the cutoffs defining mature neurons in the juvenile PL from Alderman et al. (2023). (B) Input resistance, spike duration, and threshold plotted against animal age. (C) Volcano plot of q values (FDR-corrected p values, see methods) comparing sex differences in morphological and electrical properties of adult PL neurons. Horizontal dotted line represents an alpha cutoff of 0.05.

**Extended Data Figure 3-1.**
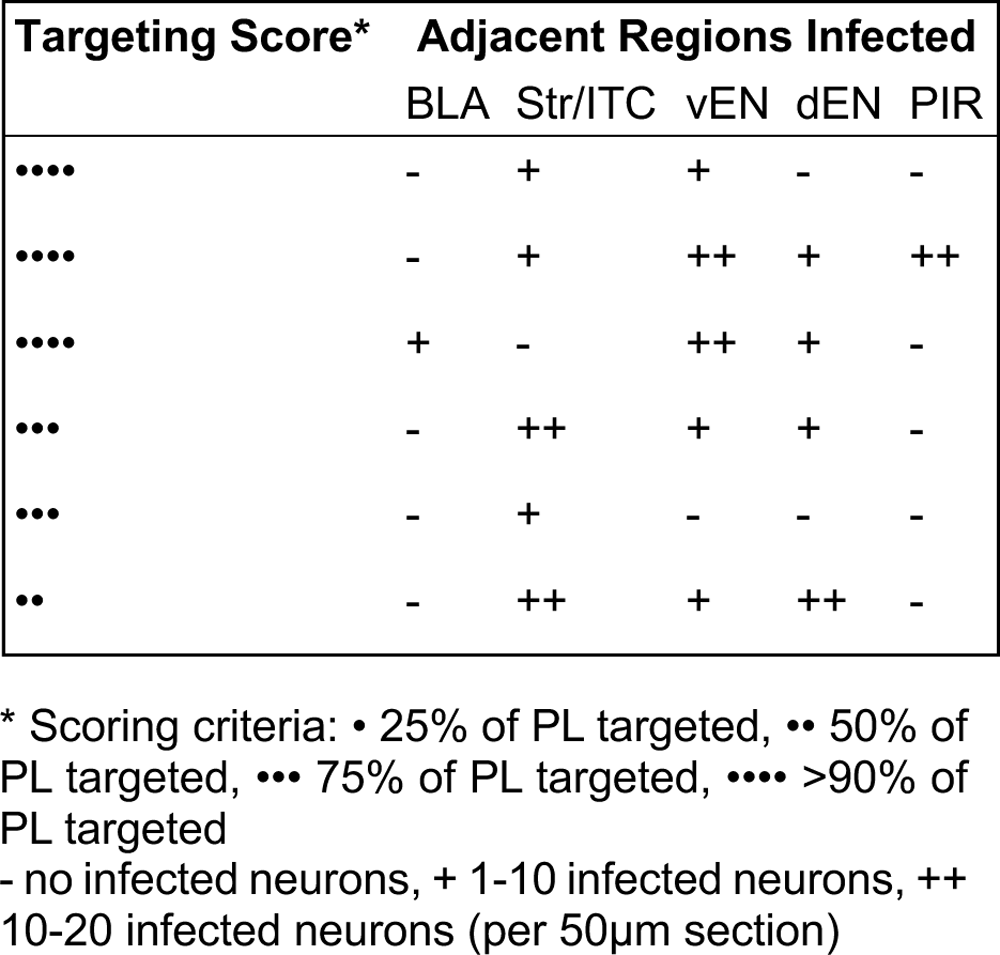
Injection targeting of retrograde virus to the adult mouse PL

**Extended Data Figure 4-1.**
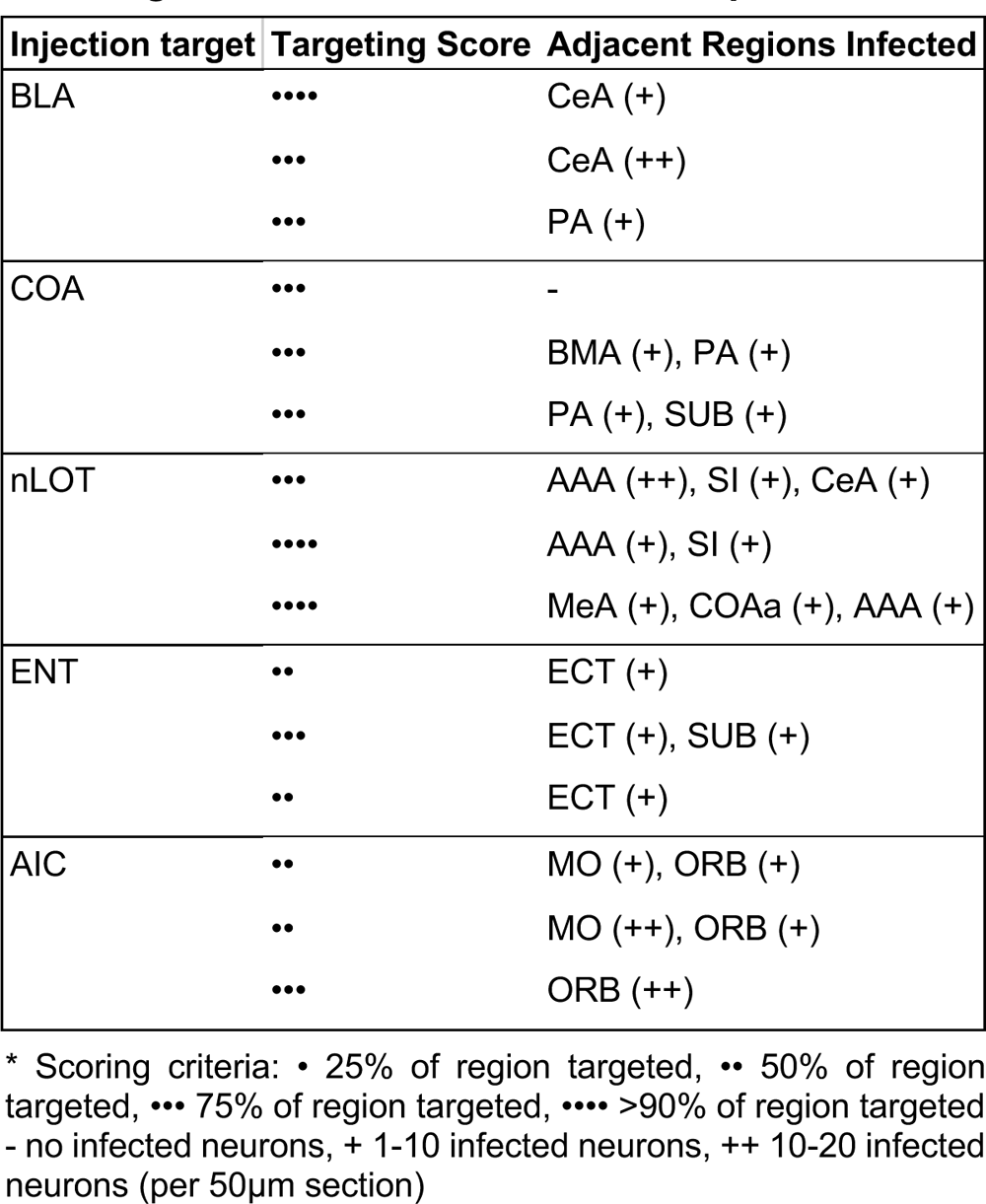
Injection targeting of anterograde virus to adult mouse PL input sources

**Extended Data Figure 5-1:**
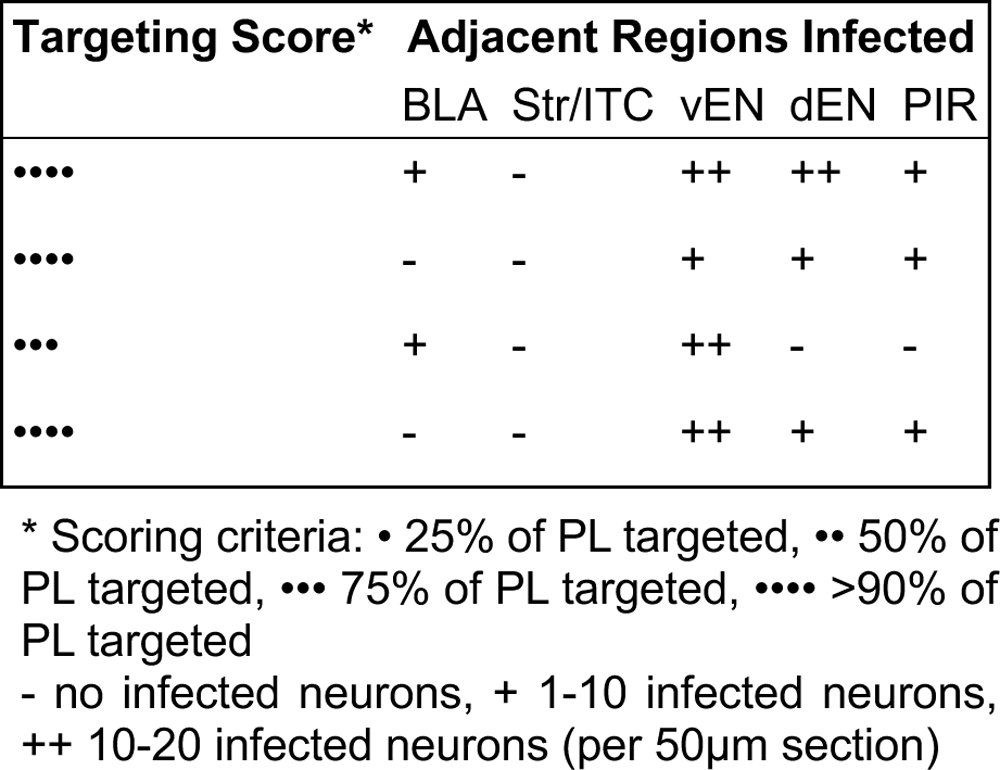
Injection targeting of anterograde virus to the adult mouse PL

## Notes

### Competing Interest Statement

The authors have declared no competing interest.

